# Insights from farming *Macrocystis pyrifera* offshore: phenotypic analysis, genome-wide association studies, genomic selection

**DOI:** 10.1101/2025.09.22.677618

**Authors:** Maxim Kovalev, Gabriel Montecinos Arismendi, Rachael M Wade, Gary Molano, Robert Miller, Daniel Reed, Filipe Alberto, Sergey Nuzhdin

**Author notes:** Corresponding author: Maxim Kovalev.

## Abstract

Seaweed farming, as a part of aquaculture, offers a sustainable alternative to modern agricultural practices; however, genetic enhancement and breeding programs for most species are underdeveloped. We aimed to advance seaweed domestication by focusing on giant kelp (*Macrocystis pyrifera*), the fastest-growing haplodiplontic brown alga, which has significant ecological and commercial importance. We analyzed phenotypic data from two offshore experimental farms conducted in 2019 and 2020, which involved hundreds of outplanted genetically diverse sporophytes. We found that outplanting season and farm design had significant effects on giant kelp biomass. Broad-sense heritability estimates showed moderate (0.27-0.50) genetic contributions to two phenotypes, carbon content and total biomass. Genome-wide association studies for these phenotypes resulted in three statistically significant SNPs, located near or within genes involved in carbohydrate metabolism and cytoskeletal functions. In addition, we applied genomic selection models that integrated sporophyte phenotypes and parental gametophyte genotypes. These models utilized reduced sets of GWAS-ranked SNPs obtained by a procedure based on linkage disequilibrium estimations. Model testing yielded cross-validation accuracy values of up to 0.84 and predictive accuracy values of up to 0.40, demonstrating the potential of marker-assisted breeding for phenotype improvement. Our results provide foundational genomic resources and tools for domesticating and breeding *M. pyrifera*, offering a basis for developing giant kelp varieties with desirable traits.

## Introduction

Modern agriculture is notorious for excessive utilization of land, water pollution, and biodiversity decline. Aside from that, climate change is causing reduced crop yields and food insecurity through alterations in weather patterns, rising temperatures, and extreme climatic events (Foley et al. 2011; Wheeler and von Braun 2013; Tilman et al. 2017). In this context, aquaculture is increasingly being viewed as an alternative food source capable of partially offsetting these issues (Gentry et al. 2017). Aquaculture is the fastest-growing food sector, producing 122.6 million tonnes of live weight worth USD 281.5 billion in 2020 (FAO 2022). Algae cultivation, dominated by marine macroalgae known as seaweeds, follows the same trend, accounting for 29% of total aquaculture production in 2020 (35.1 million tonnes) (FAO 2022). Seaweed farming is known for its many simultaneous benefits for sustainable development compared to other ocean industries. Among these are providing important source of nourishment for humans and livestock, employment opportunities that help reduce poverty, feedstocks for clean and cost-effective energy, as well as aiding climate change mitigation through innovation and responsible production (Duarte et al. 2017, 2022). Therefore, developing and improving methods of seaweed aquaculture is crucial for advancing and sustaining its benefits to society.

As of 2020, a vast majority of cultivated seaweeds were represented by just eight species, including the kelps *Saccharina japonica* (Kombu) and *Undaria pinnatifida* (Wakame). Over 99% of these seaweeds were produced in Asian countries, with China responsible for nearly 60% of the production (FAO 2022). In comparison, seaweed aquaculture is a relatively new and underdeveloped industry in the United States; however, a growing number of programs have been established to facilitate its advancement. One of these was the Macroalgae Research Inspiring Novel Energy Resources (MARINER) program funded by the Advanced Research Projects Agency-Energy (ARPA-E) of the US Department of Energy, which includes 18 innovative projects aimed at developing cost-effective methods for offshore seaweed aquaculture (Kim et al. 2019). All things considered, seaweed farming is a fast-growing sustainable industry with high potential in the US.

*Macrocystis pyrifera*, also known as giant kelp, is a brown alga (Class Phaeophyceae, Order Laminariales) with a haplodiplontic, heteromorphic life cycle, undergoing both macroscopic diploid (sporophyte) and microscopic haploid (gametophyte) phases. Giant kelp is one of the fastest-growing species on the planet, with an average elongation rate of 27 cm per day and an average biomass increase of 3.5% per day (Kain (Jones) 1982; Rassweiler et al. 2018). From an ecological perspective, *M. pyrifera* is an ecosystem engineer, and its contribution to local marine environments is an active area of research (Castorani et al. 2018; Miller et al. 2018; Lamy et al. 2020). The species is most abundant along the west coastline of the Americas (Graham et al. 2007) with the highest genetic diversity occurring in Southern California (Klingbeil et al. 2022). Among commercial applications, giant kelp is a source of alginate, as well as raw material for animal feeds, cosmetics, pharmaceuticals, fertilizers, biostimulants and biofuel (Camus et al. 2016; Chopin and Tacon 2021; Mollah et al. 2021). Wild harvesting is the primary source of *M. pyrifera* biomass, and Peru is the only country currently applying aquaculture techniques for giant kelp commercial production (Purcell-Meyerink et al. 2021). Considering economic and ecological importance, availability of genetic diversity, and seaweed farming status in the US, *M. pyrifera* stands out as an ideal candidate for domestication, cultivation, and restoration purposes.

The establishment of genetic resources and application of genetics-based selection methods can significantly facilitate and accelerate breeding programs (Jannink et al. 2010; Bhat et al. 2016; Budhlakoti et al. 2022). Nevertheless, genomic selection (GS) studies have been limited to *Saccharina latissima* (Huang et al. 2023; Bråtelund et al. 2026). Most of the genetics research for giant kelp comprised population structure investigations utilizing either microsatellite data (Alberto et al. 2010, 2011; Johansson et al. 2015) or mitochondrial DNA (Macaya and Zuccarello 2010) and a variety of transcriptomic studies using de novo transcriptome assembly (Konotchick et al. 2013; Salavarría et al. 2018; Lipinska et al. 2019; Paul et al. 2020). In 2022 and 2023, two giant kelp reference genomes were published (Paul et al. 2022; Diesel et al. 2023), with the 2023 version being the most complete and annotated, enabling fine-scale population genetics studies (Bemmels et al. 2025).

Our objective in this study was to develop genomic resources aimed at domesticating giant kelp for commercial production in Southern California. To this end, we analyzed two years (2019 and 2020) of phenotypic data of giant kelp grown at an experimental farm located in the coastal waters of Southern California. We performed genome-wide association studies (GWAS) to show that some of the observed phenotypes (total biomass and carbon content) were associated with genotype variation. We developed a genomic selection model that integrates sporophyte phenotypic and gametophyte genotypic data to evaluate the predictive power for selected traits. Furthermore, we combined this model with GWAS results to examine how subsets of associated SNPs affect the predictive power of our model.

## Methods

### Gametophyte isolation and sequencing

Obtaining gametophyte collection and performing DNA extraction and sequencing are reported in (Osborne et al. 2023) and briefly described here. First, to isolate gametophytes, reproductive blades of *Macrocystis. pyrifera* (sporophylls) were collected from four geographically and genetically different locations in Southern California in December 2018. These regions are Arroyo Quemado (AQ), Catalina Island (CI), Camp Pendleton (CP), and Leo Carrillo (LC) (**Table 1**). The sporophylls were shipped overnight to the University of Wisconsin-Milwaukee, where spore release was performed according to the Oppliger method (Oppliger et al. 2011) approximately 24 h after collection. Spores were settled onto petri dishes (in two dilutions of 10 and 100 spores per mm^2^) and placed in growth chambers (12 °C; red light, 20 μmol photons m^−2^s^−1^; 12:12 h Light:Dark photoperiod) to germinate into gametophytes with PES medium (Provasoli 1968) replaced every five weeks while the spores were in this low-density phase. Once reaching 100 μm in size, the gametophytes were individually isolated by sex and vegetatively grown under the same conditions with increased light intensity (30 μmol photons m^−2^s^−1^) to facilitate faster growth. The PES medium was replaced every week. After reaching 2 mm in diameter, clones from gametophyte cultures were selected and fragmented mechanically to promote exponential biomass growth.

**Table 1.** Description of the gametophyte collection. LD block size is defined using the threshold method (r^2^ < 0.1) with SNPs having a minor allele frequency greater than 0.05. AQ – Arroyo Quemado, CI – Catalina Island, CP – Camp Pendleton, LC – Leo Carrillo.

| Population | # of gametophytes |  |  | LD block size, Kbp |
| --- | --- | --- | --- | --- |
|  | Isolated | Sequenced | Selected/Genotyped |  |
| AQ | 66 | 64 | 63 | 32.3 |
| CI | 59 | 57 | 55 | 30.2 |
| CP | 75 | 69 | 59 | 17.9 |
| LC | 400 | 369 | 327 | 15.9 |
| *Combined* | *600* | *559* | *504* | *8.9* |

Before DNA extraction, 50-100 mg of gametophyte tissue biomass was obtained by centrifuging and discarding the supernatant for each gametophyte culture. The biomass was then pulverized using liquid nitrogen, and the NucleoSpin 96 Plant Kit (Macherey-Nagel) was used to extract high-quality genomic DNA. Library preparation and sequencing were performed by the BGI North America NGS lab (Illumina S4 Novaseq platform; 150 bp paired-end reads), resulting in 11.2 GB of 559 sequenced samples (**Table 1**).

### Sporophyte farm design and phenotyping

The full description of the farm design and sporophyte preparation can be found in Osborne et al. (2023). Here we describe the main features. Before seeding on polyvinyl lines (2 mm diameter, 6 cm long), gametophytes were fragmented into 5–10 cell-long filaments using a pellet pestle. Female gametophytes were seeded a day earlier than male ones, followed by gradual exposure to increasing white light intensities (15, 22, 35, 60 μmol photons m^−2^s^−1^) over four days. After the crosses developed into sporophytes, the polyvinyl lines were attached to nylon seeding lines (25 m long, 1/4 in diameter) at 0.5 m intervals, resulting in 50 sporophyte genotypes on each seeding line. During outplanting, divers zip-tied the seedling lines to moored long lines (1 in diameter, 137 m long). Each long line had five seeding lines with 250 sporophyte genotypes in total.

The crossing scheme for 2019 was directed to explore most of the genetic variation in limited conditions. For this purpose, 500 female gametophyte genotypes were crossed with one male gametophyte genotype of LC origin in five replicates, resulting in a total of 2,500 seeded polyvinyl lines. These polyvinyl lines were outplanted to ten moored long lines placed as five pairs of two adjacent lines, with each pair containing one replicate of the 500 unique sporophyte genotypes. The spatial positioning of the longline pairs was designed to minimize location effects on phenotypic expression (Osborne et al. 2023). Seeded lines were outplanted to the experimental farm in May 2019 and harvested for phenotypic measurements in September 2019.

The 2020 crossing design aimed to measure repeatability in the phenotypic variation observed in 2019. The number of genotypes was decreased to 96, while the number of replicates was increased to 10. The 2020 crosses were chosen based on the 2019 data. Namely, three high-biomass, low-replicate-variation females and three random females were selected from each site (24 female gametophyte genotypes in total). These females were crossed with four males, one per site, while keeping the same LC male used in 2019, resulting in 10 replicates of 96 sporophyte genotypes. These sporophytes were outplanted to four moored longlines on the experimental farm in March 2020 and harvested in July 2020.

Phenotyping of the harvested sporophytes included quantifying survivorship for each sporophyte genotype, counting the number of clonemate sporophytes growing on each polyvinyl line, total wet mass of blades, and total wet mass of blades and stipes per replicated sporophyte genotypes. Total biomass was estimated as the sum of the blades and stipes’ mass. For 2019, a portion of sporophytes was analyzed for carbon content and nitrogen content using elemental analyzer and muffle furnace combustion in the Soil and Forage Analysis laboratory at the University of Wisconsin. Populations of gametophytes, cross-populations, replicate ID, long line ID, seedling line ID, dates of outplanting and harvesting were recorded as environmental variables. The aliases for the phenotypes and environmental variables are provided in **Supplementary Table 2**.

### Analysis of genotypic data

A detailed description of sequencing quality assessment, SNP calling, and SNP processing can be found in Harden et al. (2024). Briefly, raw reads were processed to remove adapters and low-quality sequences, followed by mapping to the *M. pyrifera* nuclear genome (Diesel et al. 2023). Duplicate reads were marked, and genetic variants were called with a ploidy of 1 using GATK4 (Van der Auwera and O’Connor 2020). The resulting raw VCF file was filtered to extract biallelic SNPs, with further filtering based on scaffold presence, mean depth, and missing data percentage through vcftools (Danecek et al. 2011). The final dataset comprised 504 gametophytes (90 male and 414 female) and 1,515,399 biallelic SNPs. Missing data were imputed using an algorithm described in Harden et al. (2024), which uses LinkImpute (Money et al. 2015). SNP data were coded binary based on the absence or presence of a minor allele.

To examine genetic variation, we performed principal component analysis (PCA) and estimated the sizes of linkage disequilibrium (LD) blocks. For the former, we used the SNPRelate R package (Zheng et al. 2012) along with imputed SNP data, without any additional filtration. For the latter, we used non-imputed SNP data. Estimations were performed for each population separately and for all populations combined (five cases in total). Specifically, for each case, we used PopLDdecay (Zhang et al. 2019) on every chromosome individually, averaged the results across all chromosomes using the same software, and then fitted a monotonically decreasing curve to the averaged results. Regarding the estimations, we used 0.03, 0.05, and 0.10 minor allele frequencies (MAFs) with no restriction on the missingness frequency of the SNPs. Averaging was carried out using the parameters -bin1 10 -bin2 100 -break 1000. Fitting the decreasing curves was performed using the R package scam (Pya and Wood 2015) with the formula log10(r2) ∼ s(log10(distance), bs = “mpd”). The sizes were estimated in two ways. The first is the distance at which LD (r^2^) drops below 0.1 (threshold method). The second is the distance where LD (r^2^) is half of its maximum value (half-decay method).

### Analysis of phenotypic data

For each farm year, we constructed correlation matrices (Pearson’s correlation) exploring the phenotypes. At the same time, for the environmental variables, we used the uncertainty coefficient (UC) as a proxy for correlation since the variables are categorical. For variables *X* and *Y*, *UC*(*X*, *Y*) tells what portion of *X* can be predicted given *Y*, meaning that *UC*(*X*, *Y*) ≠ *UC*(*Y*, *X*). We focused on total biomass (Mass.Total), carbon content (Carbon), nitrogen content (Nitrogen) from the recorded phenotypes and selected cross-population (Pop.Cross), long line ID (Line.Long), seeding line ID (Line.Seed) from the environmental variables.

The association between sporophyte survivability and the selected environmental variables, including sporophyte genotypes, was tested using the Chi-squared test of independence. We separately studied how the number of clonemate sporophytes on a polyvinyl line (individual counts; Count.Ind) affected the phenotypes by organizing them into 14 groups (individual counts = 1, = 2, = 3, …, = 13, >13) and approximating trends. We then used the Kruskal-Wallis test to examine whether sporophyte genotypes, cross-population, long line ID, seeding line ID, and individual counts affect variation in the phenotypes.

Using the phenotypes affected by genotypic variation, we estimated the best linear unbiased estimations (BLUEs) representing the genetic contribution to phenotypes independent of environmental effects. To achieve this, we fitted LMM models with effects previously determined to be statistically significant by the Kruskal-Wallis test previously, using genotype as a fixed effect term and excluding cross-population, as we are directly addressing population stratification in the association studies. We also fitted these models setting genotype as a random effect, which allowed us to obtain the best linear unbiased predictors (BLUPs) for the genotype term and estimate Holland’s, the regression-based, and the shrinkage-based versions of broad-sense heritability (H^2^), defined in Schmidt et al. (2019). To fit LMM models and extract BLUEs and BLUPs, we used the R packages lme4 (Bates et al. 2015) and emmeans (Lenth 2025).

### Genome-wide association studies

To investigate associations between sporophytes’ genotypes and phenotypic data represented by BLUEs, we applied several genome-wide association models to the 2019 farm data. These models are general linear model (GLM), mixed linear model (MLM), FarmCPU (Liu et al. 2016), and Blink (Huang et al. 2019). All are implemented in the GAPIT R package (Wang and Zhang 2021). Each model was fitted with a different number of principal components (PCs; between 0 and 6) to account for population stratification. Regarding SNP data, we recreated the genotypes of the harvested sporophytes by leveraging the features of the giant kelp life cycle. Namely, since meiosis and recombination occur before spore release (Reed 1990; Le et al. 2022), sporophytes inherit one set of chromosomes from each parental gametophyte, resulting in diploid organisms with “stacked” or combined genotypes. Computationally, this means that a sporophyte’s SNP is the sum of the parental gametophytes’ SNPs when coded from the perspective of the number of minor alleles. From the genomic data, we recovered 407 and 285 sporophyte genomes that had associated BLUEs for total biomass and carbon content, respectively. For the analysis, we kept only SNPs with MAF > 0.05.

Overall, we fitted 28 GWAS models for each phenotype. We assessed the models both qualitatively and quantitatively. The former was performed by evaluating quantile-quantile (QQ) plots and the number of statistically significant SNPs based on the Benjamini-Hochberg threshold. The latter was done by estimating the genomic inflation factor (Lambda). Genes containing statistically significant SNPs or located within the LD block of those were considered putative genes affecting phenotype variations. Gene annotations (GO – Gene Ontology (Ashburner et al. 2000; The Gene Ontology Consortium et al. 2023); KOG - Eukaryotic Orthologous Groups (Tatusov et al. 1997, 2003; Galperin et al. 2025)) were captured from the *Macrocystis pyrifera* JGI genome portal (https://phycocosm.jgi.doe.gov/Macpyr2/Macpyr2.home.html). To annotate SNP effects, we used the SnpEff software (Cingolani et al. 2012).

### Genomic selection models

Genomic selection (GS) models were built using the rrBLUP R package (Endelman 2011; Endelman and Jannink 2012). The main feature of the models is that genomic estimated breeding values (GEBVs) are obtained for gametophytes based on data from sporophytes. For that, we designed an underlying LMM as follows

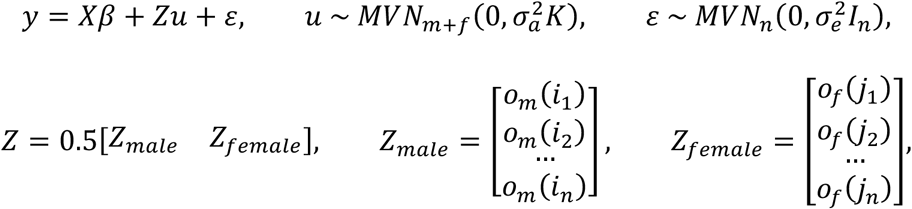

where *n* is the number of sporophytes; *m* and *f* are the total numbers of male and female gametophytes, respectively, crossed to produce the sporophytes; *y* in an *n*-vector of phenotype values; *X* is an *n* × (*p* + 1) matrix of covariates including *p* PCs and a column of 1s; *β* is a (*p* + 1)-vector of the corresponding coefficients including the intercept; *Z* is an *n* × (*m* + *f*) design block matrix represented by two parts, *Z*_*male*_ and *Z*_*female*_; *o*_*N*_(*i*) is an *N*-row where the *i*-th element equals 1, and the others are 0; *i*_1_, *i*_2_, …, *i*_*n*_ are the indexes of male gametophytes used to produce sporophyte 1,2, …, *n*, respectively; *j*_1_, *j*_2_, …, *j*_*n*_ are the indexes of female gametophytes used to produce sporophyte 1,2, …, *n*, respectively; *u* is an (*m* + *f*)-vector of random effects obeying 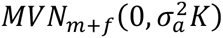, a multivariate normal distribution with a 0 mean vector and a covariance matrix 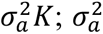 is a variance of random effects; *K* is an (*m* + *f*) × (*m* + *f*) kinship matrix; *ɛ* is an *n*-vector of errors obeying 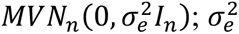 is a variance of the residual errors; *I*_*n*_ is an *n* × *n* identity matrix.

To examine the predictability of the phenotypes (BLUEs), we performed a cross-validation procedure using the 2019 data. The procedure consisted of 10,000 repetitions, where one repetition included several steps: (1) selecting 75% random sporophytes from each cross-population, (2) identifying their parental gametophytes, (3) fitting the LMM and extracting GEBVs for the parental gametophytes, (4) predicting GEBVs for the remaining gametophytes according to Clark and van der Werf (2013), (5) computing genetic additive effects (GAEs) for the remaining 25% of sporophytes as an average between male and female gametophytes’ GEBVs, (6) estimate prediction accuracy, which is Spearman correlation between the GAEs and the sporophytes’ phenotype values. Additionally, we did a separate test in which we predicted total biomass from the 2020 data using the 2019 data. For that, we identified shared sporophytes’ genotypes between the two datasets, filtered them out from the 2019 data (22 out of 407), repeated steps (2)-(4), estimated GAEs for the 2020 total biomass, and calculated Spearman correlation.

To optimize the prediction accuracy, we ran these procedures for different sets of SNPs used to obtain the kinship matrix. Specifically, for each phenotype, we performed chromosome-wide LD clumping. This involved hierarchical clustering based on LD (r^2^) between SNPs (r^2^ > 0.1) and selecting the SNP with the lowest GWAS p-value in each cluster as a representative. The two obtained sets were sorted by the p-values and used to create subsets based on ranks (for example, the top 100, top 500, top 1000, etc.). This approach was reported to be successful in multiple studies (Kaler et al. 2022; Yan et al. 2023).

## Results

### Genotypic variation in the gametophyte collection

From the spores released by the collected giant kelp sporophylls, we isolated 500 female and 100 male gametophytes. After DNA sequencing and following quality filtration, a dataset of 504 gametophytes (414 female and 90 male) with 1,515,399 biallelic SNPs was formed (**Table 1**). PCA identified four distinct genomic clusters in the PC1-PC2 plane, accounting for 5.52% of the genetic variation (**Fig. 1A**). Remarkably, these clusters matched the geographical origin of the gametophytes. In order to further examine population structure and history, we constructed LD decay curves and estimated LD block sizes. General assessment revealed increasing patterns of the sizes depending on the minor allele frequency filter for both threshold and half-decay methods (**Supplementary Fig. 1**, **Supplementary Table 1**). Specifically, for MAF > 0.05, we observed that AQ possessed the largest LD block (32.3 Kbp via the threshold method, 1200 bp via the half-decay method), while LC had the smallest one (15.9 Kbp via the threshold method, 730 bp via the half-decay method). When LD blocks were identified for the combined populations, the threshold and half-decay methods yielded 8.9 Kbp and 470 bp, respectively. LD curves for the threshold method (MAF > 0.05) are displayed in **Fig. 1B**.

**Figure 1.**
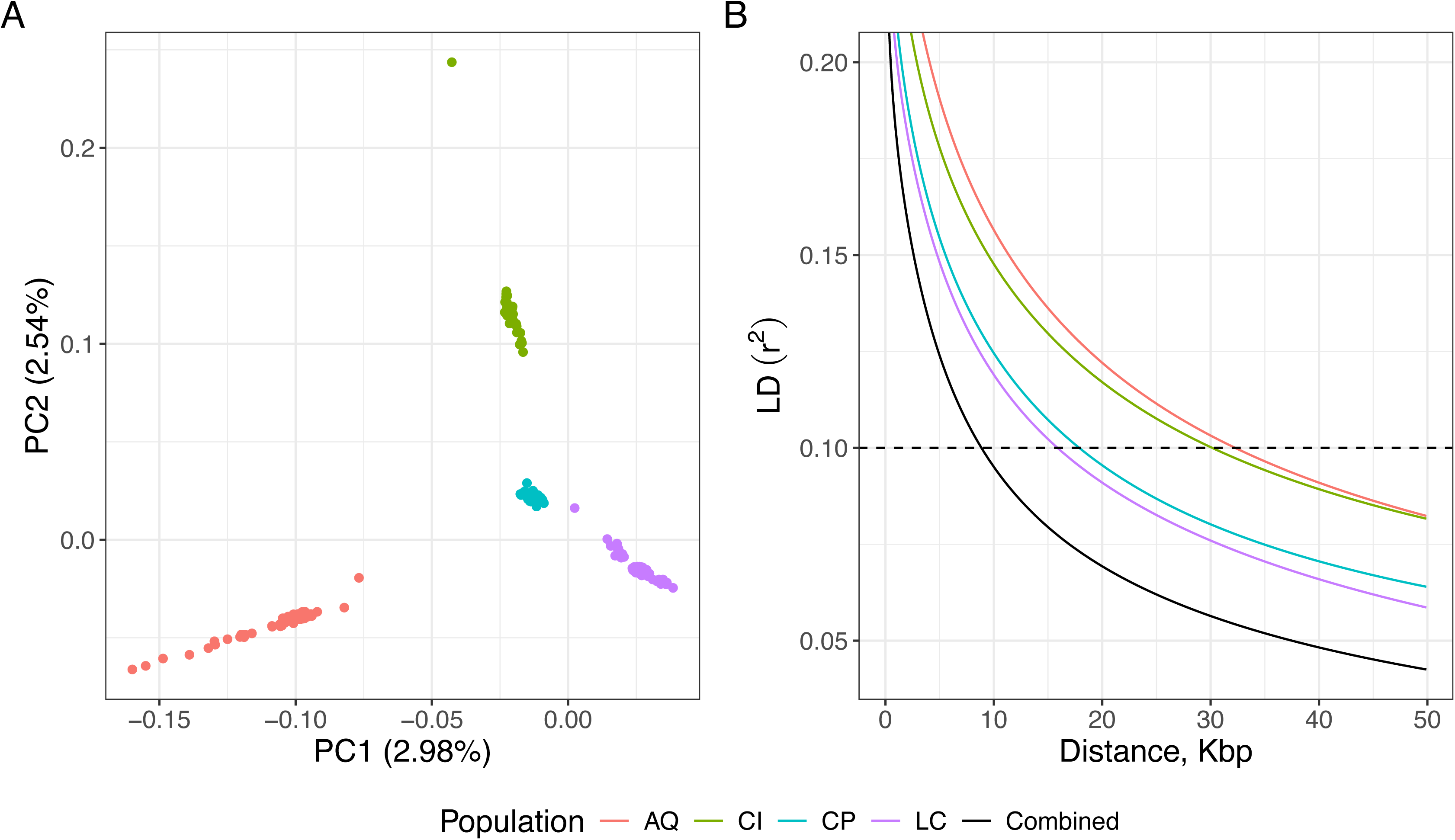
Genetic diversity of the gametophyte collection examined via (A) principal component analysis and (B) linkage disequilibrium decay.

### Description and analysis of farm data

We performed correlation analysis on the recorded phenotypes and environmental variables, the descriptions of which are provided in **Supplementary Table 2**. Cross-population (Pop.Cross) was found to be independent of other environmental variables except male and female gametophyte populations, as expected. Long line ID (Line.Long) and seeding line ID (Line.Seed) encompassed information about the remaining variables, which aligns with the non-random design of outplanting and harvesting (**Supplementary Fig. 2**). Regarding the phenotypes, all the measured mass values were positively correlated with total biomass (Mass.Total), which is their sum. In the 2019 farm, correlations among total biomass, carbon content (Carbon), and nitrogen content (Nitrogen) varied from 0.16 to 0.19, all of which were statistically significant (p ≤ 0.0001). Individual counts (Count.Ind) and total biomass were positively correlated, with Pearson’s coefficients of 0.27 and 0.55 for 2019 and 2020, respectively (p ≤ 0.0001). The correlation between individual counts and nitrogen content was 0.089 (p ≤ 0.01), whereas for carbon content, it was - 0.019 (p > 0.05) (**Supplementary Fig. 3**).

Sporophytes on 1,686 of the 2,500 lines (67.44%) outplanted in 2019, survived to be harvested, whereas sporophytes on 627 of 960 (65.31%) lines outplanted in 2020 survived to be harvested. Of those that survived, 1,669 and 614 records were analyzed for total biomass in 2019 and 2020, respectively, 1,269 records for carbon content, and 1,267 records for nitrogen content in 2019 (**Supplementary Table 3**). We used the chi-squared test of independence to analyze relationships between survivability in each year and factors such as sporophyte genotype, cross-populations, long line ID, and seeding line ID. As a result, we observed only one instance of independence, long line ID from the 2020 data (χ^2^(3, 960) = 2.484, p = 0.478), while the other cases showed statistically significant dependence (p ≤ 0.0001); all are reported in **Supplementary Table 4**.

In this study, we aimed to analyze three phenotypes: total biomass, carbon content, and nitrogen content. Their distributions are shown in **Fig. 2**. Total biomass (wet mass) ranged between 1 and 1,002 g in 2019 and between 1 and 4,414 g in 2020. Carbon content constituted between 18.05 and 26.32 % of dry mass, while the nitrogen content ranged between 0.78 and 1.99 % of dry mass. Comparing total biomass between the two years, the median for 2019 was 110 g, whereas the median for 2020 was 405 g.

**Figure 2.**
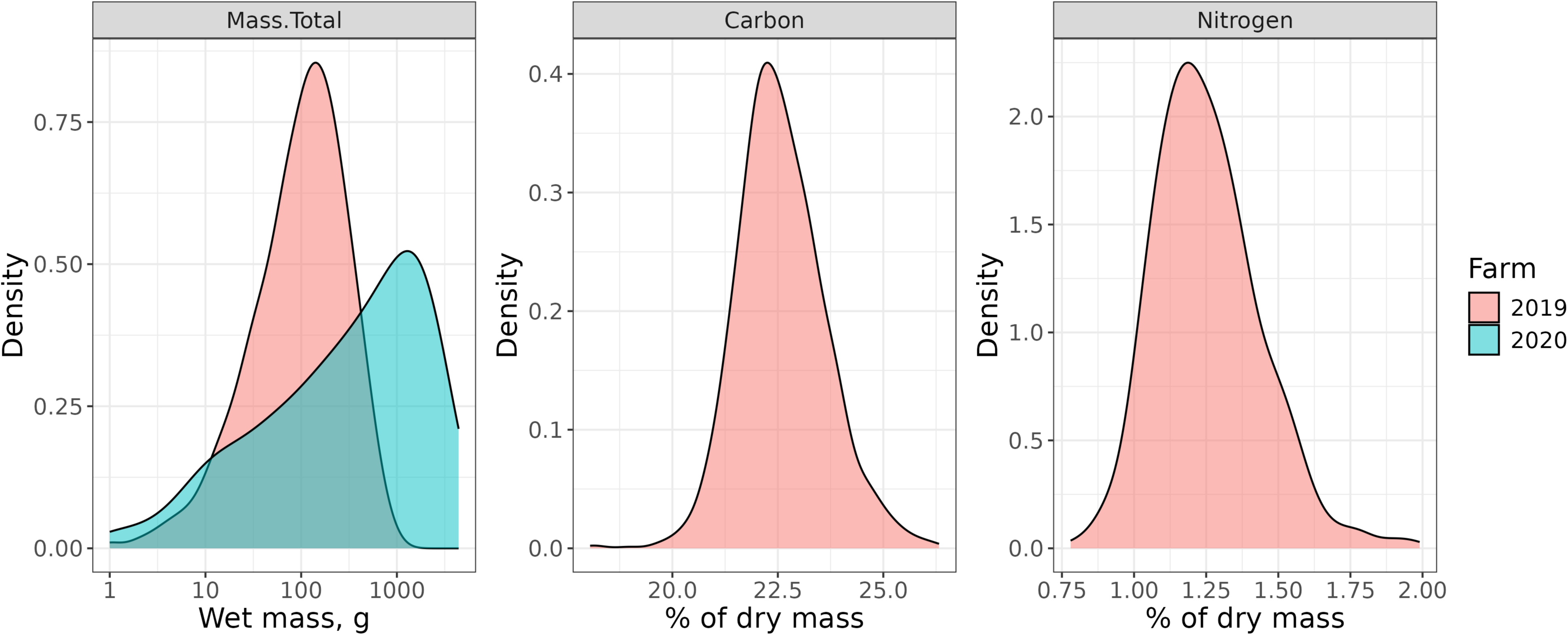
Distributions of the phenotypes from 2019 and 2020 farm data. Mass.Total – total biomass, Carbon – carbon content, Nitrogen – nitrogen content.

Twenty-four genotypes were shared between the two years. We performed non-parametric two-way ANOVA (Wobbrock et al. 2011; Kay et al. 2025) to analyze the effect of genotype and year on total biomass. The analysis revealed that there was no statistically significant interaction between the genotype and year effects (F(23, 233) = 1.4237, p = 0.1). However, the main effects of genotype and year were statistically significant (p ≤ 0.001). The total biomass values for these 24 genotypes were averaged and illustrated in **Fig. 3**, representing the difference between averaged biomass distributions across the two years.

**Figure 3.**
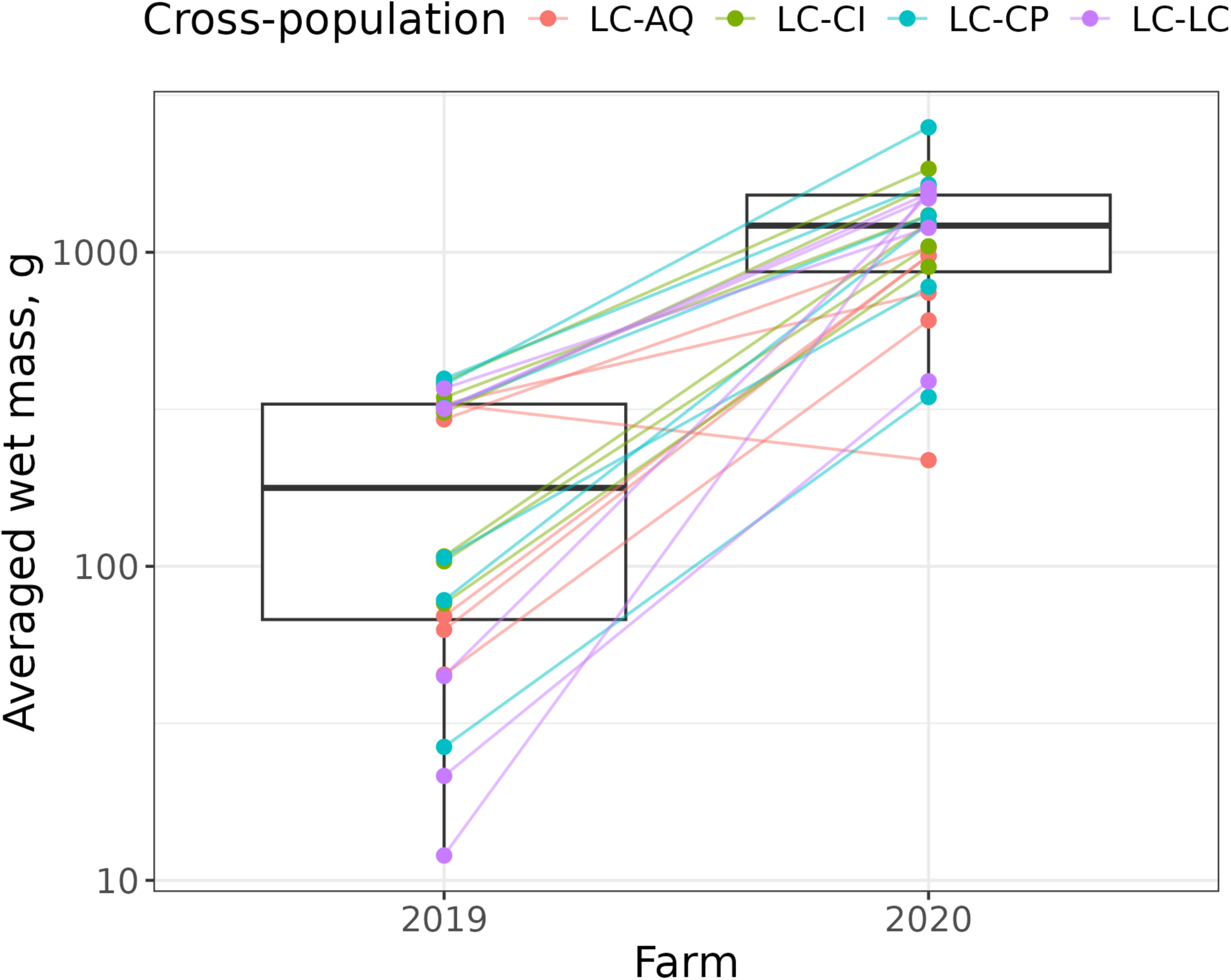
Performance comparison of the same crosses (n = 24) between the two farms.

We were interested in investigating the relationship between the number of clonemate sporophytes on 6 cm long polyvinyl lines, which we refer as individual counts (Count.Ind), and total biomass on the lines at the time of harvest. For this, we organized the former into 14 groups (Count.Ind = 1,= 2, = 3, …, = 13, >13) and evaluated the distributions of total biomass in these groups (**Supplementary Fig. 4A**). Qualitatively, the estimated trends showed similar relationships for smaller individual counts. The difference became more pronounced for counts greater than five. In 2019, the trend remained unchanged on average, but decreased when the individual counts exceeded 13. This contrasts with the 2020 trend, where an increase in individual counts leads to a monotonic increase in total biomass. Similarly, we visualized the effect of individual counts on carbon content and nitrogen content (**Supplementary Fig. 4B**).

To assess the effect of individual counts, as well as the effects of genotype, cross-population, long line ID, and seeding line ID on the phenotypes quantitatively, we utilized a Kruskal-Wallis test (**Supplementary Table 5**). All the mentioned factors demonstrated statistical significance in both years, except for three cases. The effect of individual counts on carbon content was not statistically significant (H(13) = 12.58, p = 0.481). Likewise, the effects of genotype (H(344) = 341.60, p = 0.526) and cross-population (H(3) = 7.55, p = 0.056) on nitrogen content were insignificant. Based on these results, we excluded nitrogen content from the downstream analysis.

For the subsequent steps, we obtained best linear unbiased estimates (BLUEs) and best linear unbiased predictions (BLUPs) for total biomass (n = 491 and 96 for 2019 and 2020, respectively) and carbon content (n = 345). The distributions of BLUEs per cross-population are displayed in **Supplementary Fig. 5**. Calculating both BLUEs and BLUPs allowed us to perform different estimations of broad-sense heritability (**Supplementary Table 6**). For total biomass, broad-sense heritability estimates ranged from 0.27 to 0.28 in 2019 and from 0.47 to 0.50 in 2020. For carbon content, broad-sense heritability varied between 0.31 and 0.35.

### Genome-wide association studies for carbon content and total biomass

We screened 378,063 and 379,690 SNPs for carbon content and total biomass, respectively. Out of the total 56 GWAS models, only one model yielded significant results for the former and three for the latter phenotypes (**Supplementary Table 7**). For total biomass, we chose the Blink model, as it had the genetic inflation factor value closest to one. The results of these models are represented by Manhattan and QQ plots in **Fig. 4**. We were able to identify two SNPs associated with carbon content and one SNP associated with total biomass (**Table 2**).

**Figure 4.**
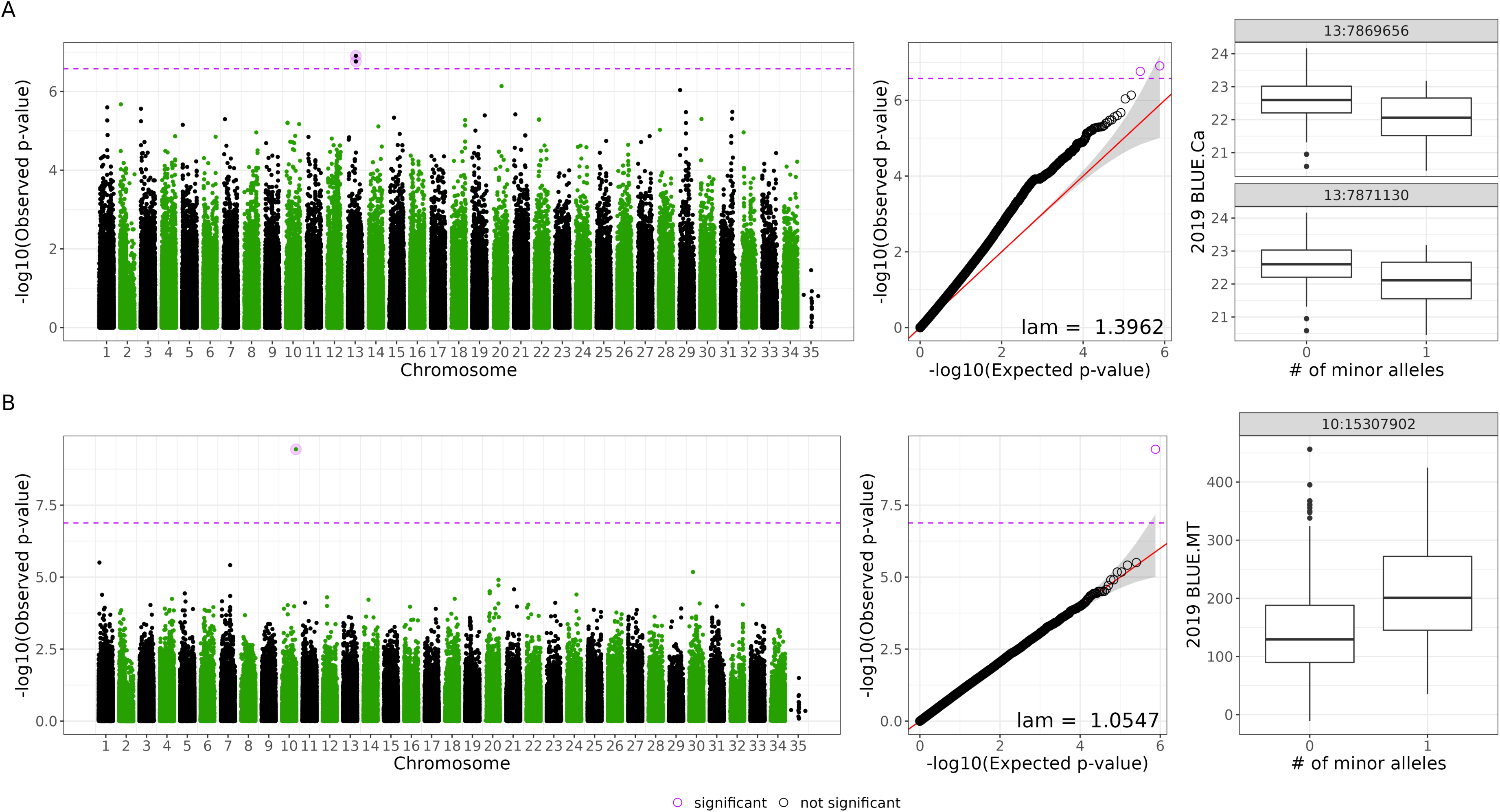
Manhattan plot, quantile-quantile (QQ) plots, and BLUE distributions for (A) carbon content and (B) total biomass.

**Table 2.**
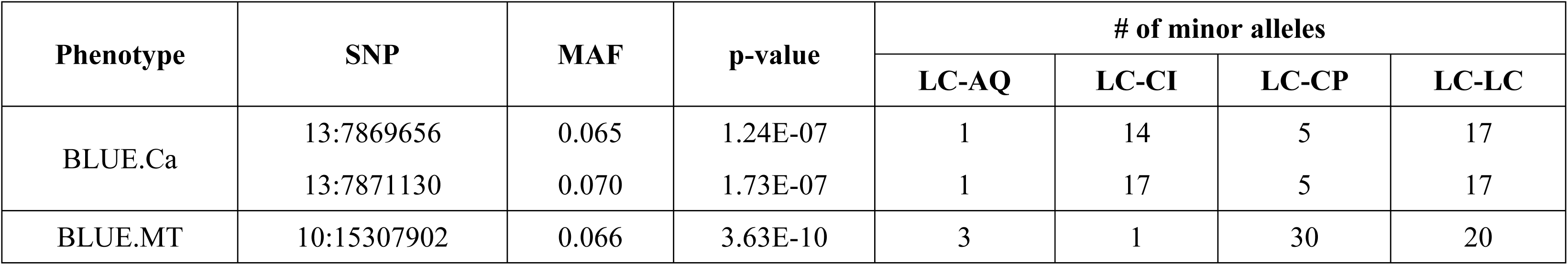
Summary of the statistically significant SNPs and cross-population contributions to total minor allele numbers. BLUE.Ca – carbon content BLUEs; BLUE.MT – total biomass BLUEs; MAF – minor allele frequency.

Carbon content seemed to be affected negatively by the presence of minor alleles in the identified SNPs, opposite to the total biomass case (**Fig. 4**). These “negative effect minor alleles” tended to be more abundant in sporophytes from LC-CI and LC-LC cross-populations. The distributions of minor alleles associated with total biomass were slightly different. The majority of them were in sporophytes from LC-CP and LC-LC cross-populations. The range of MAF for all three identified SNPs was between 0.065 and 0.070.

Using the LD block size estimated for all populations combined (8.9 Kbp), we identified four relevant genes in total, jgi.p|Macpyr2|9543804 and jgi.p|Macpyr2|9808315 for the carbon content SNPs, jgi.p|Macpyr2|9800916 and jgi.p|Macpyr2|9800917 for the total biomass SNPs (**Table 3**). According to the annotations, gene products of jgi.p|Macpyr2|9543804, jgi.p|Macpyr2|9808315, and jgi.p|Macpyr2|9800916 were predicted to be glutathione S-transferase, fructose-bisphosphate aldolase, and dynein heavy chain, respectively. For carbon content, the identified genes are involved in carbohydrate transport and metabolism, as well as posttranslational modification. For total biomass, we observed only one annotated gene, whose function was associated with the cytoskeleton. The SNP 13:7869656, belonging to a coding region of the jgi.p|Macpyr2|9543804 gene, was annotated as a missense variant by SnpEff, leading to an amino acid substitution from threonine to alanine at position 45 of the protein.

**Table 3.** Summary and annotations of genes located within the LD window of the statistically significant SNPs. BLUE.Ca – carbon content BLUEs, BLUE.MT – total biomass BLUEs, GO – Gene Ontology database, KOG – Eukaryotic Orthologous Groups database.

| Phenotype | Gene | Related SNPs | SNPs' location | Annotations |  |  |  |  |  |
| --- | --- | --- | --- | --- | --- | --- | --- | --- | --- |
|  |  |  |  | KOG (Description) | KOG (Class) | KOG (Group) | GO (Cellular component) | GO (Biological process) | GO (Molecular function) |
| BLUE.Ca | jgi.p\|Macpyr2\|9543804 | 13:7869656 | Coding region | (KOG2903)<br>Predicted glutathione S-transferase | Posttranslational modification, protein turnover, chaperones | CELLULAR PROCESSES AND SIGNALING | - | - | - |
|  |  | 13:7871130 | Intronic |  |  |  |  |  |  |
|  | jgi.p\|Macpyr2\|9808315 | 13:7869656 | Intergenic (1039 bp upstream) | (KOG1557)<br>Fructose-biphosphate aldolase | Carbohydrate transport and metabolism | METABOLISM | - | (GO:0006096)<br>glycolysis | (GO:0004332) fructose-bisphosphate aldolase activity |
|  |  | 13:7871130 | Intergenic (2513 bp upstream) |  |  |  |  |  |  |
| BLUE.MT | jgi.p\|Macpyr2\|9800916 | 10:1530790<br>2 | Intronic | (KOG3595) Dyneins, heavy chain | Cytoskeleton | CELLULAR PROCESSES AND SIGNALING | (GO:0030286)<br>dynein complex | (GO:0007018)<br>microtubule-based movement | (GO:0000166) nucleotide binding; (GO:0003777) microtubule motor activity; (GO:0005524) ATP binding; (GO:0016887) ATPase activity; (GO:0017111) nucleoside-triphosphatase activity |
|  | jgi.p\|Macpyr2\|9800917 | 10:1530790<br>2 | Intergenic (5850 bp downstream) | - | - | - | - | - | - |

### Genomic selection models

Identification of LD-based representative SNPs within chromosomes based on their association with the phenotypes resulted in 94,412 SNPs for carbon content and 94,393 SNPs for total biomass. Using the representative SNPs, we assessed the potential of genomic selection through cross-validation on the 2019 data, as well as tested the genomic selection models on data they were not trained on (2020 and 2019 data, respectively). Results are illustrated in **Fig. 5**.

**Figure 5.**
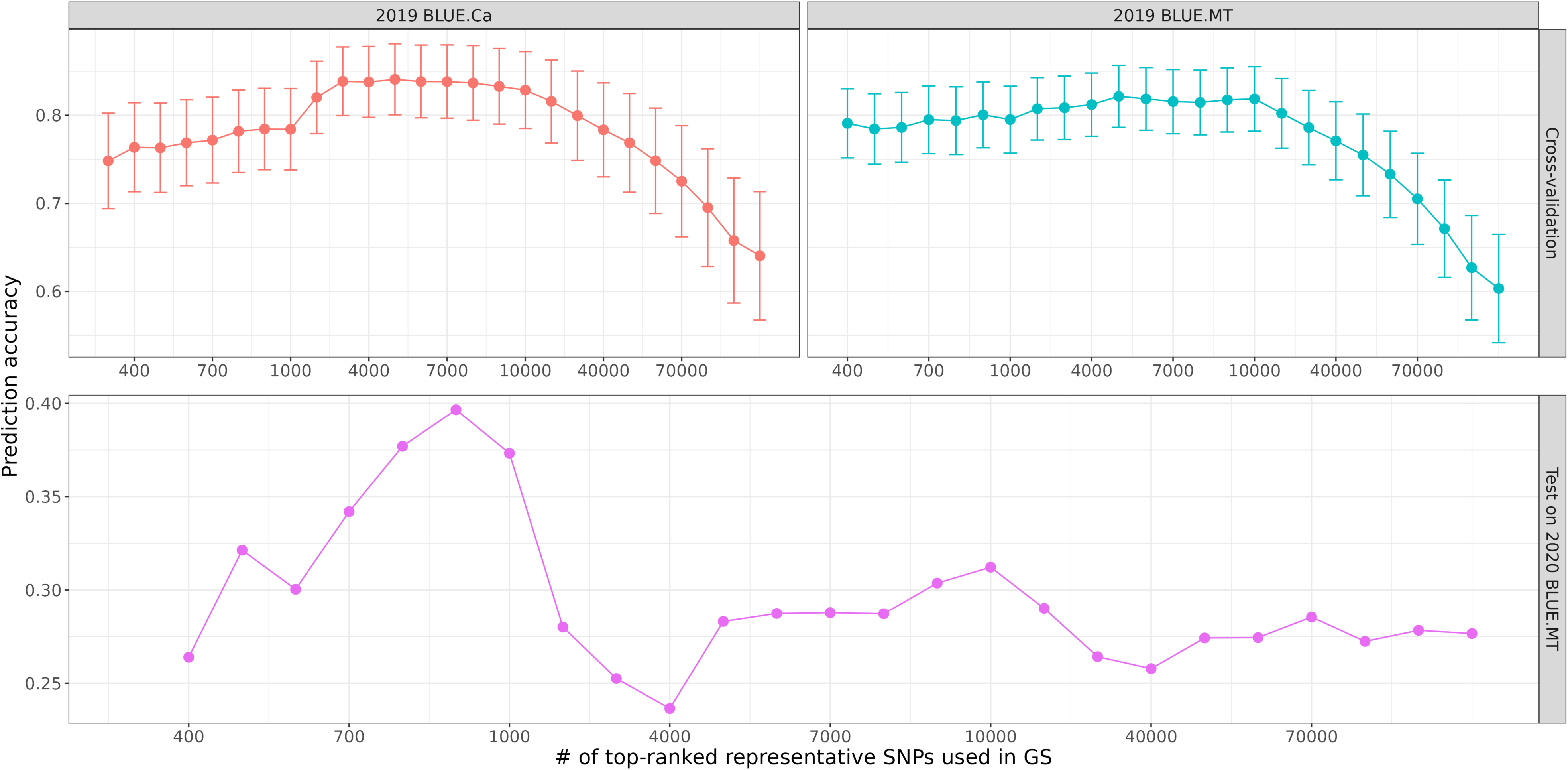
Prediction accuracies (Spearman correlation) for genomic selection (GS) models based on different numbers of GWAS-ranked representative SNPs used in the kinship matrix estimation. The top two panels depict the cross-validation procedure results (mean ± sd) obtained for the 2019 data. The bottom panel illustrates GS models trained on 2019 total biomass BLUEs and tested on the 2020 ones. BLUE.Ca – carbon content BLUEs, BLUE.MT – total biomass BLUEs; the rightmost point for each trend reflects the case when all independent SNPs are used.

In the case of cross-validation, the trends in prediction accuracy for phenotypes were similar when plotted against the number of GWAS-ranked SNPs used. Accuracy initially increased to a certain point, followed by mild fluctuations, and then declined rapidly after using 10,000 representative SNPs in the model. For both phenotypes, the highest mean accuracy was achieved at 5,000 top-ranked representative SNPs, with values of 0.84 for carbon content and 0.82 for total biomass. When actually testing genomic selection models, prediction accuracy was within the 0.23-0.40 interval, with the highest accuracy of 0.396 at 900 top-ranked representative SNPs. The accuracy trend was increasing until 900 SNPs, followed by a decline and fluctuations around 0.275. Remarkably, when all representative SNPs were used, the prediction accuracy value was 0.277.

## Discussion

### Population structure and linkage disequilibrium

Principal component analysis identified four well-supported genetic clusters that exactly matched the source locations (AQ, CI, CP, LC). These clusters demonstrated a clear spatial structure consistent with previous observations of dispersal capacity and environmental variation in southern California giant kelp populations (Johansson et al. 2015). It is remarkable that not a single individual was misclassified in terms of population placement. This suggests extremely low levels of concurrent kelp migration rates, despite heavy ocean traffic between populations. However, actual estimations require dense sampling and the identity-by-descent analysis.

Across the entire collection, comprising 504 genotyped gametophytes, the LD block size calculated via the threshold method (MAF > 0.05) extended only to 8.9 Kbp while population-specific values varied substantially, with the largest in AQ (32.3 Kbp) and the smallest in LC (15.9 Kbp). Assessing LD patterns between populations, LC and CP showed the fastest LD decay, followed by CI and AQ, indicating differences in their recombination rates and effective population sizes. These results also align with the microsatellite allelic richness differences among the populations described in (Johansson et al. 2015).

For comparison, Diesel et al. (2023) analyzed 48 diploid macroscopic sporophytes of giant kelp from Santa Barbara, Camp Pendleton, and Catalina Island. To obtain LD block sizes, they applied the threshold method with MAF > 0.10. Reported sizes were only 5.5–6 Kbp, which is 10 times smaller compared to our estimates acquired in the same way (**Supplementary Table 1**). The discrepancy in LD block sizes between sporophytes and gametophytes indicates higher levels of LD in the latter. Interestingly, this result would be expected for broadcast-spawner species with sweepstakes reproductive success (SRS) (Hedgecock and Pudovkin 2011). Giant kelp has at times been categorized as a broadcast spawner (Bringloe et al. 2020); it has a haplodiplontic life cycle, undergoing meiosis in specialized tissues in the diploid sporophyte stage to release millions of haploid microscopic zoospores (Leal et al. 2014), which differentiate into gamete-producing gametophytes. The reproductive success of giant kelp is known to depend on a high number of factors, such as spore settlement density (less than 1 mm apart for successful fertilization), light, substrate, nutrients, and water motion (Molano et al. 2022). Nevertheless, giant kelp population genetics has not been tested under the SRS model, and additional simulation studies are required to investigate this.

A study of the Asian kelp *Undaria pinnatifida* (Graf et al. 2021) provided LD block size estimates for 41 sporophytes. Calculations were performed using the half-decay method and varied between 3.14 Kbp and 27.33 Kbp. Even though a minimal allele frequency of 0.0365 was used instead of the minor allele frequency during the filtration step, our results for all MAFs we used ranged between 0.4 and 2.4 Kbp across populations and between 0.19 and 1.2 Kbp when the populations were combined.

### Field performance of farmed crosses

About one-third of out-planted sporophytes failed to survive at both experiments, while genotypic loss was minimal (1.8% in 2019 and 0% in 2020; **Supplementary Table 3**), indicating that most mortality was caused by environmental factors rather than genetic reasons. This is further supported by the chi-squared test results (**Supplementary Table 4**), where the lowest p-values were associated with variability among long lines and seedling lines. Interestingly, the variation attributed to these sources was greater for the 2019 experiment which occurred between May and September. We believe our mortality rates are within a normal range. To the best of our knowledge, there are no reported mortality rates specifically from kelp offshore farms; however, there are studies documenting mortality rates for other cases. For comparison, reports of wave-induced mortality in natural populations of giant kelp ranged between 2% and 94% depending on depth and year of observation (Seymour et al. 1989). From a marine heatwave study, average mortality rates of *Saccharina latissima* (sugar kelp) historically ranged from 33% to 49% in summer in the eastern USA (Filbee-Dexter et al. 2020). For sugar kelp restoration efforts, mortality rates between March and September ranged between 12.5% and 50% across multiple years (Sogn Andersen et al. 2011).

A general comparison of total biomass distributions from the two farm experiments revealed pronounced differences, with the 2020 farm exhibiting higher biomass values. When shared crosses between the farms were compared, non-parametric two-way ANOVA showed that the only significant effect was the year of outplanting. Assuming that the main difference between the farms was the timeline of outplanting (2019 summer vs. 2020 spring), these results indicate that yield is highly dependent on the time of year outplanting is done. Similar results were reported for giant kelp farmed in Southeast Alaska (Stekoll et al. 2021). Sporophytes outplanted in February-March tended to have larger plant length compared to those outplanted in April-May. Our broad-sense heritability estimations also indicate that the environmental impact on total biomass was higher in 2019.

Testing how our phenotype variances were affected by different factors revealed several key findings. First, all phenotypes except carbon content were significantly dependent on the number of individual sporophytes in replicates (**Supplementary Table 5**, **Supplementary Fig. 3**). Interestingly, the total biomass dependency varied with the year of outplanting (**Supplementary Fig. 4A**). Compared to the monotonically increasing pattern in the 2020 farm, we observed a plateau in total biomass after the number of clonemates reached six, followed by a decrease when the number approached 13. We suspect that variations in outplanting season and resource competition influenced the differences between the observed trends. Nutrient availability is known to be lower and temperatures are higher during summer compared to winter in Southern California (Zimmerman and Kremer 1984). In the competition context, space and light are viewed as essential resources that influence both interspecific and intraspecific competition at every stage of the *M. pyrifera* life cycle (Graham et al. 2007). Secondly, among all recorded phenotypes, only nitrogen content was not statistically affected by the genotype or cross-population of the crosses (**Supplementary Table 5**). This aligns with the concept that nitrogen content is highly dependent on the availability of nutrients, as demonstrated in various experimental studies (Gerard and North 1980; Gerard 1982; Zimmerman and Kremer 1986). Additionally, it has been stated that during periods of limited nutrient availability, kelp increasingly utilizes regenerated forms of nitrogen, such as ammonium and urea (Lees et al. 2024). Evidence suggests that the kelp microbiome plays an important role in supplying its host with nitrogen in its most readily assimilated form (Florez et al. 2021; Hochroth and Pfister 2024). However, our data do not suffice to fully support these statements.

### Genotype-phenotype association for carbon content and total biomass

Genome-wide association studies for our traits resulted in three statistically significant SNPs in total, two from chromosome 13 for carbon content and one from chromosome 10 for total biomass. Notably, all the models that produced statistically significant results were not corrected for population stratification (via including principal components as covariates). Minor alleles for the SNPs were mostly concentrated in two cross-populations (LC-CI for carbon content, LC-CP for total biomass, and LC-LC for both). These findings suggest that our associations may be confounded with the population structure. This is further supported by assessing the relationship between the traits and the principal components (**Supplementary Fig. 6**).

Nevertheless, the biological signal we observed is relevant to the context of this study. The SNPs associated with carbon content lie within an ∼9 kb LD block containing fructose-1,6-bisphosphate aldolase (FBA) and glutathione S-transferase (GST) genes. FBA is a key enzyme involved in both catabolic (glycolysis) and anabolic (gluconeogenesis and Calvin–Benson cycle) pathways (Yang et al. 2023). Its importance was tested in multiple studies. Overexpression of plastid FBA in tobacco plants enhanced photosynthesis and plant growth under a constant light intensity and a high CO_2_ concentration (Uematsu et al. 2012). In tomatoes, overexpressing SlFBA4, one of the chloroplast/plastid FBA, enhanced their photosynthetic capability and protected cell membranes from chilling stress (Cai et al. 2022). Carbon assimilation and metabolism in rice were also reported to be affected when the expression of OsFBA1 is manipulated (Liu et al. 2025). GST, a protein superfamily with diverse functions, is involved in many plant processes, including detoxification of xenobiotics, secondary metabolism, growth and development, and protection against both biotic and abiotic stresses (Vaish et al. 2020). Although there is no direct link to carbon metabolism, studies suggest GST can play an intermediate role by altering the carbon metabolite pool (Mueller et al. 2000; Sappl et al. 2009). For both identified SNPs, minor alleles shift the distributions of the phenotype so that the median carbon content becomes lower. Moreover, one SNP (13:7869656) was defined as a missense variant in the GST gene by SnpEff, causing a p.Thr45Ala substitution. However, additional analysis is required to understand the morphological and metabolic consequences of this substitution.

We identified only one statistically significant SNP for total biomass, which was located within an intronic region of a dynein heavy chain (DHC) gene, and it also fell within the LD block overlapping with a gene that has no annotation. DHCs are cytoskeletal motor proteins that perform various functions, such as cargo transport along cytoplasmic microtubules, retrograde transport in the cilia and flagella, and contributions to spindle assembly and positioning during mitosis, to name but a few (Roberts et al. 2013; Hou and Witman 2015; Hinchcliffe and Vaughan 2018). We hypothesize that involvement in mitosis and transport functions can be connected to growth rates, and thereby, to biomass accumulation (Goranov et al. 2009; Weraduwage et al. 2015). The presence of the minor allele in the identified variant is associated with an increase in biomass. To confirm the association between the gene and total biomass, additional experimental steps, such as CRISPR/Cas9 or RNAi knockdown, are required.

Given the presence of the population structure, we conducted an additional GWAS analysis for the largest cross-population group – LC-LC (sample size is 270 for total biomass and 180 for carbon content). The use of common SNPs (MAF > 0.05) did not yield any statistically significant results. Including more rare variants (0.01 < MAF ≤ 0.05) led to four models with the results for carbon content. The best model (Lambda ≍ 0.984) identified six significant SNPs associated with four genes, of which only two had annotations. Those are Ankyrin (KOG4177) and Amino acid transporter protein (KOG1305). To the best of our knowledge, previous research has not established a connection between the associated genes and carbon metabolism or assimilation.

### Efficiency of genomic predictions

After clustering SNPs within chromosomes based on their LD and selecting a representative SNP with the lowest p-value in each cluster, the total number of SNPs was reduced to approximately 94,000 for both traits. This approach allowed us to minimize redundancy due to local linkage while keeping relatively associated SNPs. Using this pruned set, we observed that both cross-validation and actual predictions followed a similar pattern. Namely, prediction accuracy increased with the number of top-ranked SNPs, peaked or plateaued, and then declined rapidly as additional SNPs were included, which is consistent with a similar approach applied to data in other taxa (Heinrich et al. 2023).

In our models trained on 2019 data, cross-validation for the 2019 crosses consistently achieved higher accuracy than phenotype prediction for the 2020 crosses. This was expected, as the cross-validation used training and testing subsets from the same dataset. The peak prediction accuracy was observed at around 5,000 SNPs for the cross-validation, whereas it was approximately 900 SNPs for the actual predictions. These results can be explained via the curse of dimensionality (CoD) and overfitting. The CoD reduces the effectiveness of distance-based models, including genomic selection, as an increase in the number of dimensions (SNPs) leads to data sparsity, causing distances between points to become less informative (Altman and Krzywinski 2018). In other words, the more SNPs we include to calculate the kinship matrix, the less meaningful relatedness values with respect to a trait in question. From the cross-validation point of view, a model deals with only one dataset – one distribution of data. As a result, we favor more complex models (with more SNPs used) because we evaluate performance within the same distribution, even if that complexity leads to overfitting. By contrast, when trying to identify optimal SNPs by training a model on one dataset and testing on another, we impose generalization ability on the model and avoid overfitting.

### Implications for giant kelp aquaculture in the Southern California Bight

This study has been designed to enable genetics-based breeding programs for giant kelp. Throughout our analyses, we observed that there is a decent degree of heritable variation for key production phenotypes, such as biomass and carbon content, within and between kelp populations. However, and consistent with (Johansson et al. 2015), we also observe strong population subdivision and significant phenotypic differences between populations, which align with strong GWAS signals being confounded by population structure. For faster breeding progress, we propose starting with crosses between populations to unlock the most helpful genetic variation. Unfortunately, this conflicts with regulatory limitations, which require only local kelps to be used for local farming and do not permit the production of any F2 individuals from interpopulation crosses. Overall, the potential for converting kelps into very efficient, robust, and uniform crops appears very high; however, additional strategies, such as careful control of domesticated kelp reproduction (Vissers et al. 2024), must be considered as a part of any breeding program for ocean crops of the future.

## Acknowledgements

We would like to thank Christie Yorke, Clint Nelson, and all other team members from the University of California, Santa Barbara, the University of Wisconsin-Milwaukee, and the University of Southern California who contributed to the harvest and phenotyping of the kelp farms, and the NSF grant (OCE-1831937) for financial support. We also like to thank University of Southern California researchers Kelly DeWeese, Bernadeth Tolentino, and Kyle Allen for their valuable comments and discussions during manuscript preparation.

This work is funded by Macroalgae Research Inspiring Novel Energy Resources at the Advanced Research Projects Agency—Energy at the Department of Energy (DE-AR-0000914) and NOAA National Sea Grant Program (through the Early Propagation Strategies for Aquaculture Species call)

## Author contribution statement

Isolation, maintenance, and propagation of gametophytes: GMA, FA; DNA extractions and sequences processing: GMA, GM, MK; producing sporophytes from gametophyte crosses: GMA; harvest and phenotyping of sporophytes: DR, RM, GMA, GM; data processing: MK, RMW, FA; phenotypic analysis, association studies, and breeding models: MK; interpretation of results: MK, GM, RMW; manuscript preparation: MK, RMW, GM, DR, RM, FA, SN; study concept and design: SN, FA, DR, RM.

## Data archiving

The raw Illumina sequences of 559 gametophytes used to generate the SNP data are available in the NCBI database under the accession code PRJNA1050779 (https://www.ncbi.nlm.nih.gov/sra/PRJNA1050779). Other data and sources supporting the findings are available from the corresponding author upon reasonable request.

