## Supplementary materials for "Insights from farming *Macrocystis pyrifera* offshore: phenotypic analysis, genome-wide association studies, genomic selection"

### Supplementary Figures

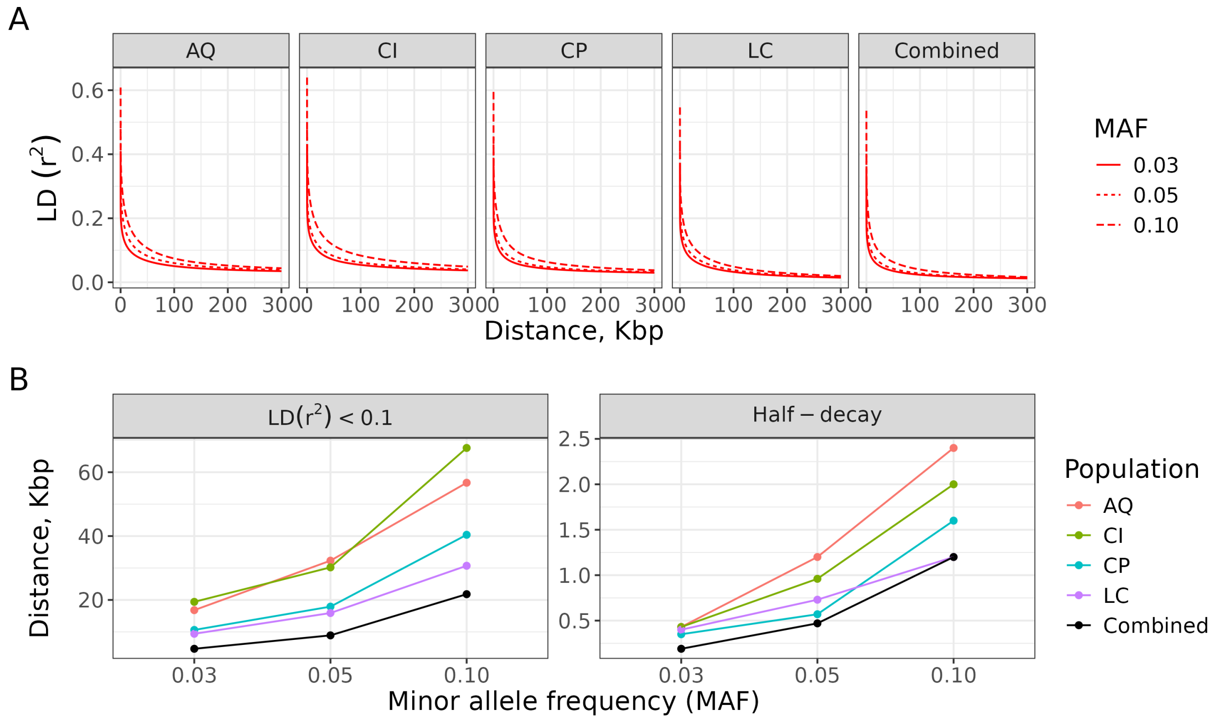


**Supplementary Figure 1.** LD analysis of the gametophyte collection. (A) – LD decay curves for the populations separated and combined when the SNP data have different minor allele frequency filtration. (B) – LD block sizes estimated via the threshold method (left panel) and the half-decay method (right panel). AQ – Arroyo Quemado, CI – Catalina Island, CP – Camp Pendleton, LC – Leo Carrillo.

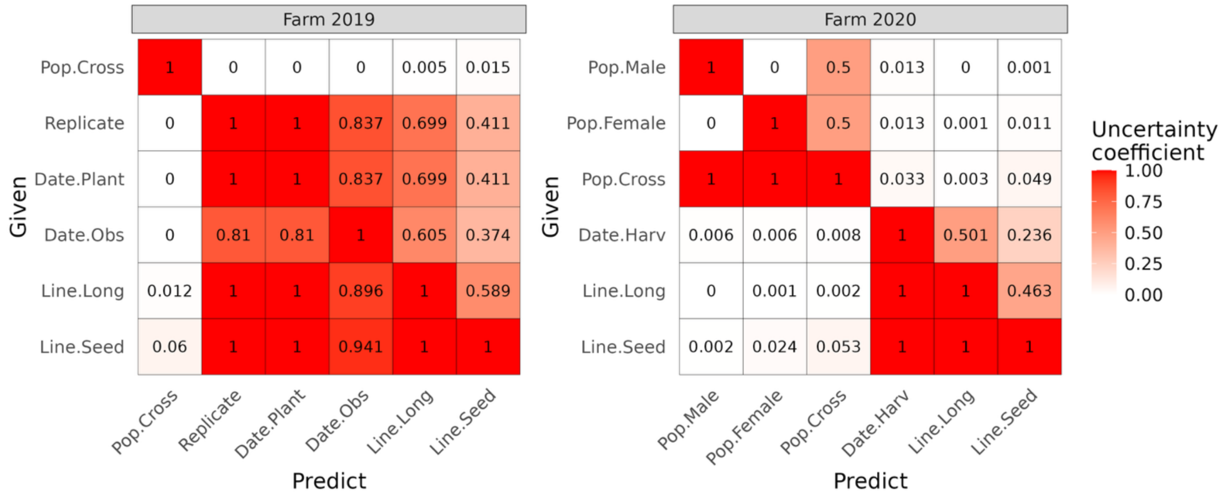


**Supplementary Figure 2.** Analysis of the environmental variables recorded for the 2019 and 2020 farms.

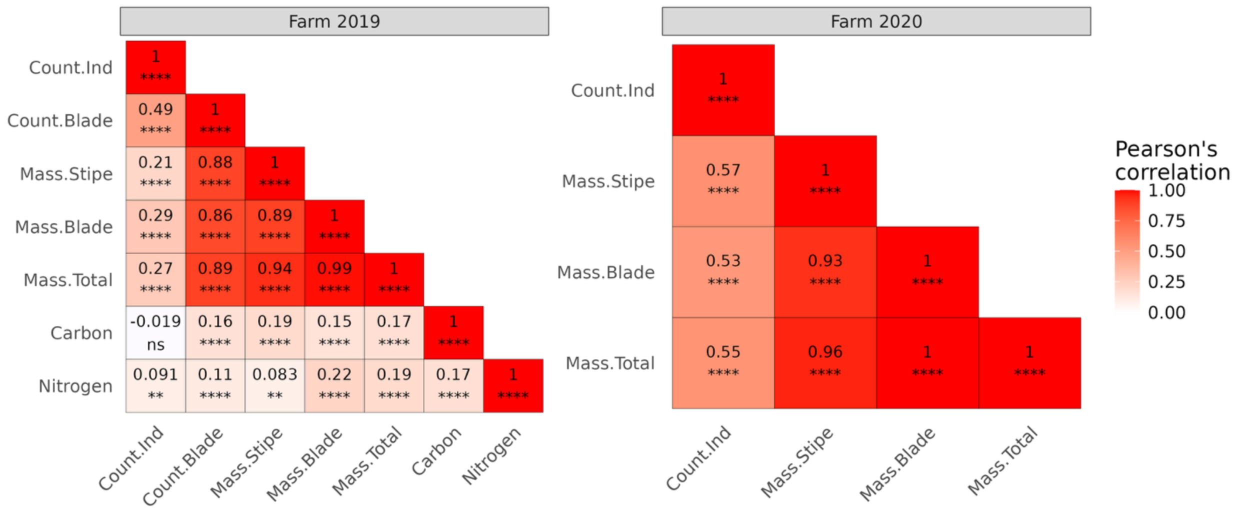


**Supplementary Figure 3.** Correlation analysis of the phenotypes measured for the 2019 and 2020 farms. ns – not significant, * – p-value ≤ 0.05, ** – p-value ≤ 0.01, *** – p-value ≤ 0.001, **** – p-value ≤ 0.0001.

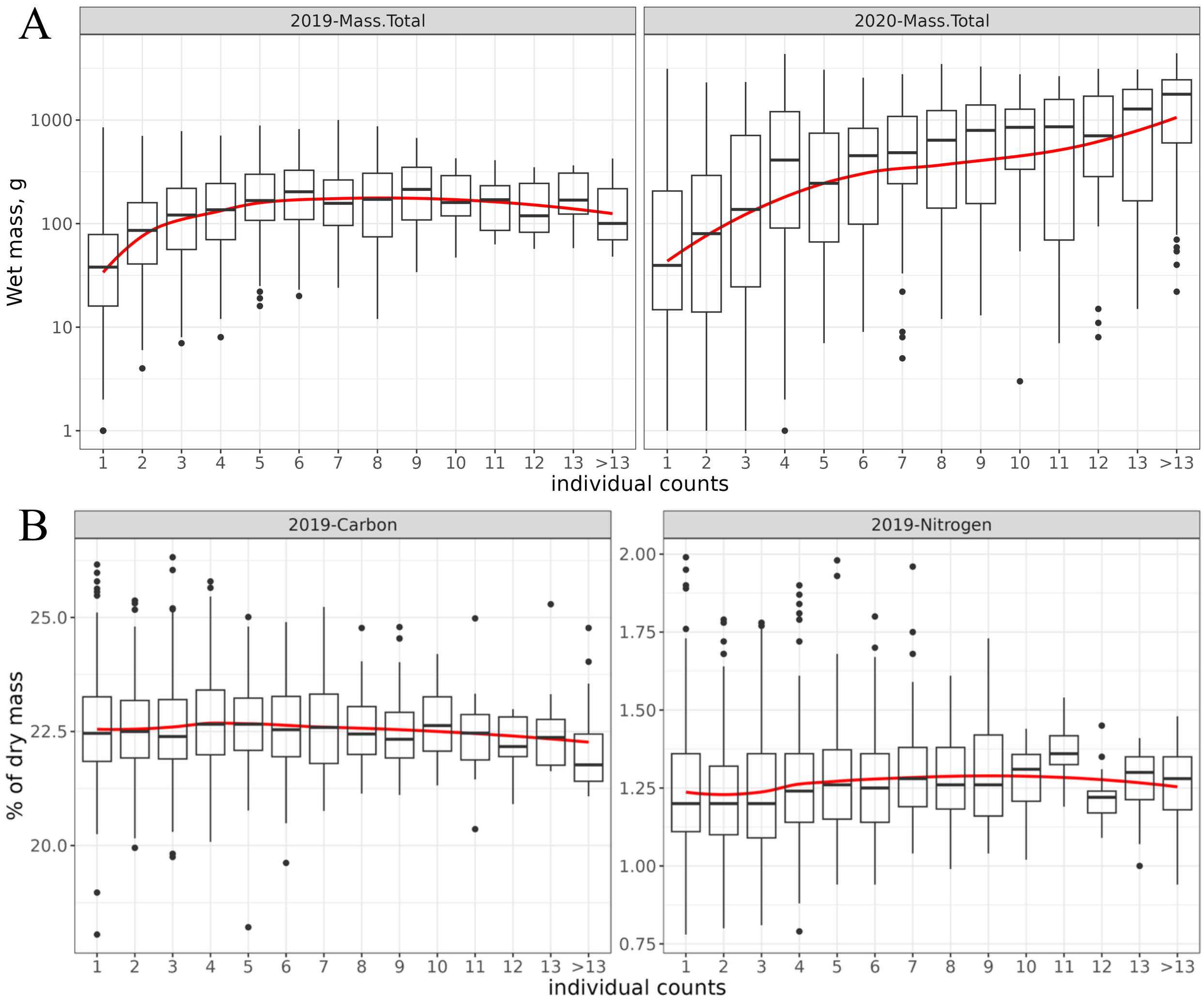


**Supplementary Figure 4.** Distributions of (A) total biomass, (B) carbon content, and nitrogen content relative to individual counts (the number of clonemate sporophytes on a polyvinyl line). The red line shows the trends of the phenotypes over individual counts formed as groups.

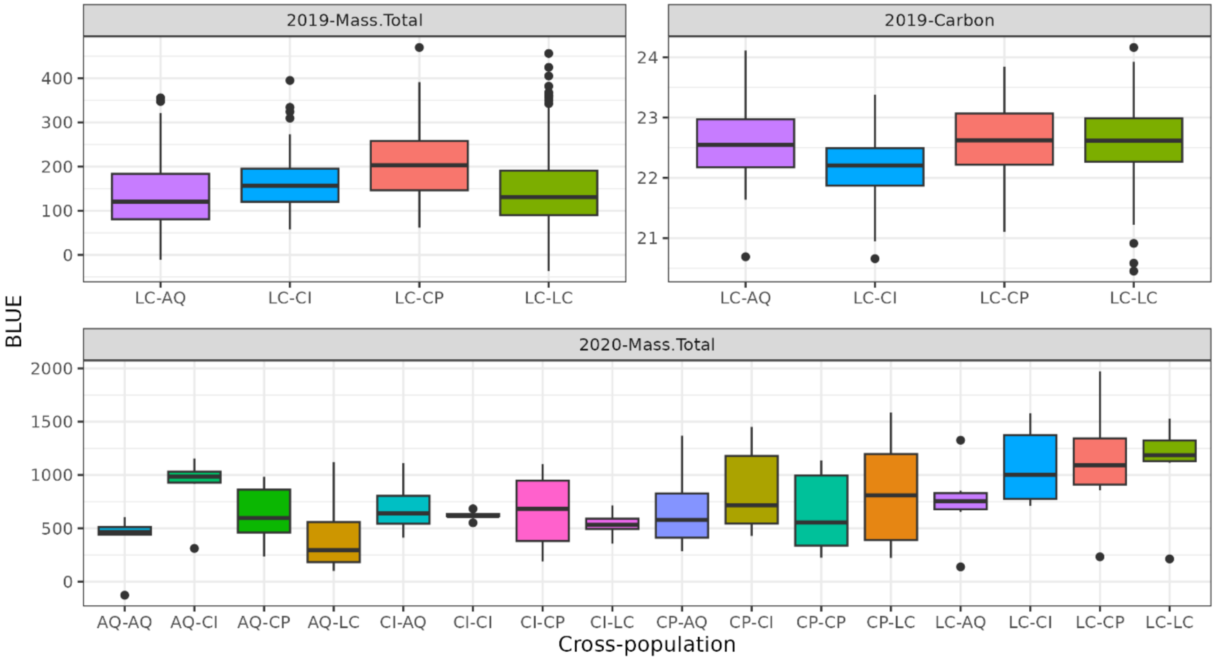


**Supplementary Figure 5.** Distributions of the phenotypes' BLUEs relative to cross-population.

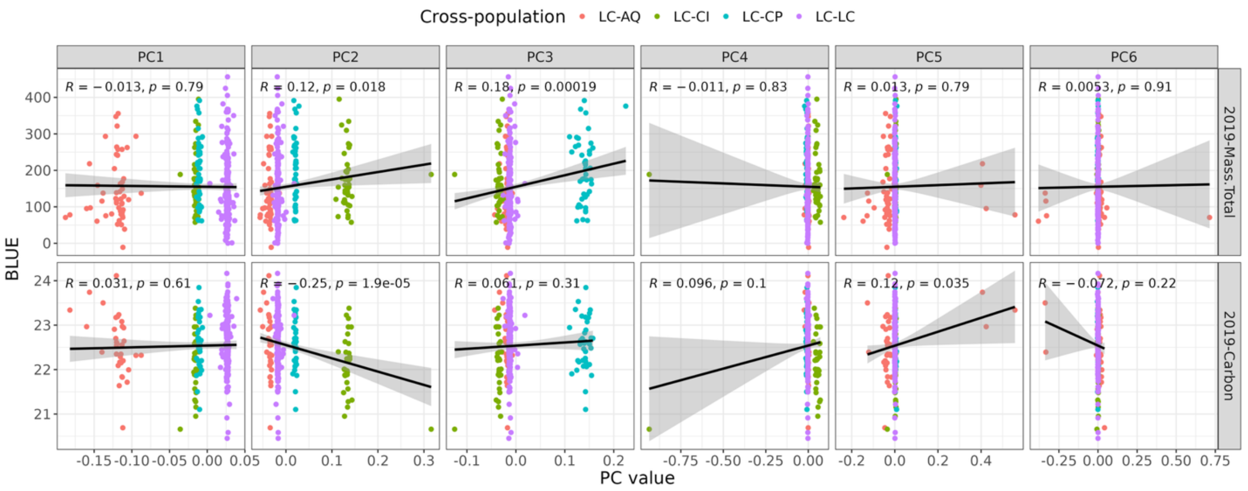


**Supplementary Figure 6.** Relationships between principal components and BLUEs for 2019 total biomass and carbon content

### Supplementary Tables

| Population | Threshold ($r^2 < 0.1$) | | | Half-decay | | |
|---|---|---|---|---|---|---|
| | **MAF > 0.03** | **MAF > 0.05** | **MAF > 0.10** | **MAF > 0.03** | **MAF > 0.05** | **MAF > 0.10** |
| AQ | 16.8 | 32.3 | 56.7 | 0.43 | 1.2 | 2.4 |
| CI | 19.4 | 30.2 | 67.6 | 0.43 | 0.96 | 2 |
| CP | 10.6 | 17.9 | 40.4 | 0.35 | 0.57 | 1.6 |
| LC | 9.4 | 15.9 | 30.7 | 0.4 | 0.73 | 1.2 |
| ***Combined*** | 4.7 | 8.9 | 21.8 | 0.19 | 0.47 | 1.2 |

**Supplementary Table 1.** LD block sizes estimated via the threshold ($r^2 < 0.1$) and half-decay methods for different minor allele frequency filters. The sizes are given in Kbp. MAF – minor allele frequency, AQ – Arroyo Quemado, CI – Catalina Island, CP – Camp Pendleton, LC – Leo Carrillo.

| | | | |
|---|---|---|---|
| Phenotypes | Common | Count.Ind | Number of clonemates (individual sporophytes per replicate on a polyvinyl string) |
| | | Mass.Stipe | Wet mass in grams of stipes and pneumatocysts (no blades or holdfasts) from all clonemates in a replicate |
| | | Mass.Blade | Wet mass in grams of blades from all clonemates in a replicate |
| | | Mass.Total | Sum of stipe and blade biomass in grams |
| | 2019 | Count.Blade | Number of blades counted |
| | | Carbon | Carbon content, % of dry mass |
| | | Nitrogen | Nitrogen content, % of dry mass |
| Variables | Common | Pop.Male | Population of male gametophyte |
| | | Pop.Female | Population of female gametophyte |
| | | Pop.Cross | Cross-population; formed as Pop.Male-Pop.Female (for example, LC-AQ) |
| | | Date.Plant | Date sporophytes were outplanted |
| | | Date.Harv; Date.Obs | Date sporophytes were harvested (for 2019 phenotypes were observed at the same day) |
| | | Line.Long | Long line ID |
| | | Line.Seed | Seeding line ID |
| | 2019 | Replicate | The replicate ID of a cross |

**Supplementary Table 2.** Description of the phenotypes and environmental variables.

| Year | Phenotype | Total | | Survived | | Survived, % | | Phenotyped | | Phenotyped, % | |
|---|---|---|---|---|---|---|---|---|---|---|---|
| | | **Spo** | **Gen** | **Spo** | **Gen** | **Spo** | **Gen** | **Spo** | **Gen** | **Spo** | **Gen** |
| 2019 | Mass.Total | 2500 | 500 | 1686 | 491 | 67.44 | 98.2 | 1669 | 491 | 66.76 | 98.2 |
| | Carbon | | | | | | | 1269 | 345 | 50.76 | 69 |
| | Nitrogen | | | | | | | 1267 | 345 | 50.68 | 69 |
| 2020 | Mass.Total | 960 | 96 | 627 | 96 | 65.31 | 100 | 614 | 96 | 63.96 | 100 |

**Supplementary Table 3.** Description of the phenotypes' dynamics from outplanting to phenotyping. Spo – sporophytes, Gen – genotypes.

| Variable | Farm | N | Df | $\chi 2$-statistic | p-value |
|---|---|---|---|---|---|
| Genotype | 2019 | 2500 | 499 | 654.70 | 3.19E-06 |
| | 2020 | 960 | 95 | 178.28 | 5.01E-07 |
| Pop.Cross | 2019 | 2500 | 3 | 32.54 | 4.03E-07 |
| | 2020 | 960 | 15 | 58.89 | 3.92E-07 |
| Line.Long | 2019 | 2500 | 9 | 337.59 | 2.71E-67 |
| | 2020 | 960 | 3 | 2.48 | ***4.78E-01*** |
| Line.Seed | 2019 | 2500 | 49 | 408.05 | 4.20E-58 |
| | 2020 | 960 | 19 | 51.89 | 6.87E-05 |

**Supplementary Table 4.** Chi-square tests of independence between sporophytes' survivability and selected environmental variables, including genotypes. N – number of datapoints, Df – degree of freedom; bold italic font illustrates insignificant results (p-value > 0.05).

| Phenotype | Farm | Variable | N | Df | H-statistic | p-value |
|---|---|---|---|---|---|---|
| Mass.Total | 2019 | Genotype | 1669 | 490 | 741.39 | 1.45E-12 |
| | | Pop.Cross | | 3 | 32.70 | 3.73E-07 |
| | | Line.Long | | 9 | 67.78 | 4.15E-11 |
| | | Line.Seed | | 49 | 113.85 | 4.47E-07 |
| | | CI.Group | | 13 | 426.21 | 6.45E-83 |
| | 2020 | Genotype | 614 | 95 | 180.43 | 2.95E-07 |
| | | Pop.Cross | | 15 | 70.67 | 3.39E-09 |
| | | Line.Long | | 3 | 16.99 | 7.10E-04 |
| | | Line.Seed | | 19 | 102.36 | 2.00E-13 |
| | | CI.Group | | 13 | 180.53 | 1.32E-31 |
| Carbon | 2019 | Genotype | 1269 | 344 | 430.34 | 1.05E-03 |
| | | Pop.Cross | | 3 | 26.58 | 7.21E-06 |
| | | Line.Long | | 9 | 97.31 | 5.49E-17 |
| | | Line.Seed | | 49 | 182.21 | 3.24E-17 |
| | | CI.Group | | 13 | 12.58 | ***4.81E-01*** |
| Nitrogen | 2019 | Genotype | 1267 | 344 | 341.60 | ***5.26E-01*** |
| | | Pop.Cross | | 3 | 7.55 | ***5.62E-02*** |
| | | Line.Long | | 9 | 130.82 | 8.05E-24 |
| | | Line.Seed | | 49 | 204.11 | 7.90E-21 |
| | | CI.Group | | 13 | 35.45 | 7.23E-04 |

**Supplementary Table 5.** Testing the influence of sporophyte genotypes and specific environmental variables on phenotypes' variance using the Kruskal-Wallis test. N – number of datapoints, Df – degree of freedom; bold italic font illustrates insignificant results (p-value > 0.05).

| **Phenotype** | **Farm** | **Broad-sense heritability ($H^2$)** | | |
|---|---|---|---|---|
| | | **Holland's** | **Regression-based** | **Shrinkage-based** |
| Mass.Total | 2019 | 0.282 | 0.270 | 0.270 |
| | 2020 | 0.471 | 0.496 | 0.488 |
| Carbon | 2019 | 0.309 | 0.341 | 0.348 |

**Supplementary Table 6.** Broad-sense heritability estimations for the selected phenotypes.

| Phenotype | Model | # of used PCs | # of significant SNPs | Lambda | Rank |
|---|---|---|---|---|---|
| BLUE.Ca | GLM | 0 | 2 | 1.3962 | 1 |
| BLUE.MT | Blink | 0 | 1 | 1.0547 | 1 |
| | FarmCPU | 0 | 12 | 0.9497 | 2 |
| | GLM | 0 | 1 | 1.4342 | 3 |

**Supplementary Table 7.** Summary of the successful GWAS models ranked for each phenotype using Lambda. BLUE.Ca – carbon content BLUEs, BLUE.MT – total biomass BLUEs, Lambda – genetic inflation factor.
